# CORACLE (COVID-19 liteRAture CompiLEr): A platform for efficient tracking and extraction of SARS-CoV-2 and COVID-19 literature, with examples from post-COVID with respiratory involvement

**DOI:** 10.1101/2024.03.18.584627

**Authors:** Kristina Piontkovskaya, Yulian Luo, Pia Lindberg, Jing Gao, Michael Runold, Iryna Kolosenko, Chuan-Xing Li, Åsa M. Wheelock

## Abstract

**Background:** During the COVID-19 pandemic there emerged a need to efficiently monitor and process large volumes of scientific literature on the subject. Currently, as the pandemic is winding down, the clinicians encountered a novel syndrome - Post-acute Sequelae of COVID- 19 (PASC) - that affects over 10% of those who contract SARS-CoV-2 and presents a significant and growing challenge in the medical field. The continuous influx of new research publications underscores a critical need for efficient tools for navigating the literature.

**Objectives:** We aimed to develop an application which will allow monitoring and categorizing COVID-19-related literature through building publication networks and medical subject headings (MeSH) maps to be able to quickly identify key publications and publication networks.

**Methods:** We introduce CORACLE (COVID-19 liteRAture CompiLEr), an innovative web application designed for the analysis of COVID-19-related scientific articles and the identification of research trends. CORACLE features three primary interfaces: The “Search” interface, which displays research trends and citation links; the “Citation Map” interface, allowing users to create tailored citation networks from PubMed Identifiers (PMIDs) to uncover common references among selected articles; and the “MeSH” interface, highlighting current MeSH trends and associations between MeSH terms.

**Results:** Our web application, CORACLE, leverages regularly updated PubMed data to aggregate and categorize the extensive literature on COVID-19 and PASC, aiding in the identification of relevant research publication hubs. Using lung function in PASC patients as a search example, we demonstrate how to identify and visualize the interactions between the relevant publications.

**Conclusion:** CORACLE proves to be an effective tool for the extraction and analysis of literature. Its functionalities, including the MeSH trends and customizable citation mapping, facilitate the discovery of relevant information and emerging trends in COVID-19 and PASC research.

## The COVID-19 crisis underscores the need for more evolved literature mining tools

The emergence of the severe acute respiratory syndrome coronavirus 2 (SARS-CoV-2) pandemic, and the associated respiratory coronavirus disease (COVID-19), has undeniably posed one of the most significant threats to global public health in recent history, impacting populations worldwide. In response to this unprecedented crisis, a collective effort among clinicians, researchers, and scientific communities has fostered a remarkable effort in open-source dissemination of research findings. This endeavor aimed to expedite the resolution of the global health crisis by accelerating scientific progress, as evidenced by the large volume of literature dedicated to SARS-CoV-2 and COVID-19-related topics.

The proliferation of COVID-19 literature is striking, with over 400,000 papers published to date. Even as the pandemic is currently winding down, an influx of 212 new publications was recorded in the past week alone (week 11 of 2024). Additionally, a substantial body of literature addresses previous coronavirus outbreaks, including those caused by SARS-CoV and MERS-CoV, which have afflicted populations with severe respiratory syndromes over the past decade. Furthermore, the emergence of post-acute COVID-19 syndrome in the aftermath of the COVID-19 pandemic also prompted a substantial growing number of publications that demand daily monitoring for the involved professionals. The exponential growth of COVID-19-related publications presents a challenge in staying updated on the status of the field. To facilitate access to relevant research, several open-access literature repositories have been established, such as LitCovid ^1^, the WHO COVID-19 Database ^2^, and the Coronavirus Knowledge Hub ^3^. These hubs offer comprehensive collections of publications, featuring search and filtering functionalities akin to those found in databases like PubMed. However, navigating through the vast array of publications necessitates considerable human curation, requiring significant time investment and domain expertise. Given the dynamic nature of the field and the rapid influx of new research, there is a pressing need for refined and time-efficient literature mining tools. To address this demand, we present CORACLE (COvid liteRAture CompiLEr), a literature integration tool tailored to support researchers, clinicians, epidemiologists, policy-makers, and other involved personnel in navigating, tracking, filtering, summarizing, and prioritizing COVID-related literature. CORACLE aims to provide daily updates on the latest trends and findings in the COVID field, offering a valuable resource to aid in their pursuit of knowledge related to COVID-19 and post-COVID-19 syndrome (https://coracle.cmm.se).

### Structure and function of the CORACLE workflow

The objective of the CORACLE application is to deliver a fast, simple, and smooth user experience when navigating large numbers of COVID-19-related publications. It is a web-based application and does not require any installation locally on the users’ computers. To provide a smooth user experience, this application follows the structure of a Single Page Application (SPA) with back-end and front-end separated. **Figure 1A** shows a schematic overview of the project.

**Figure 1.**
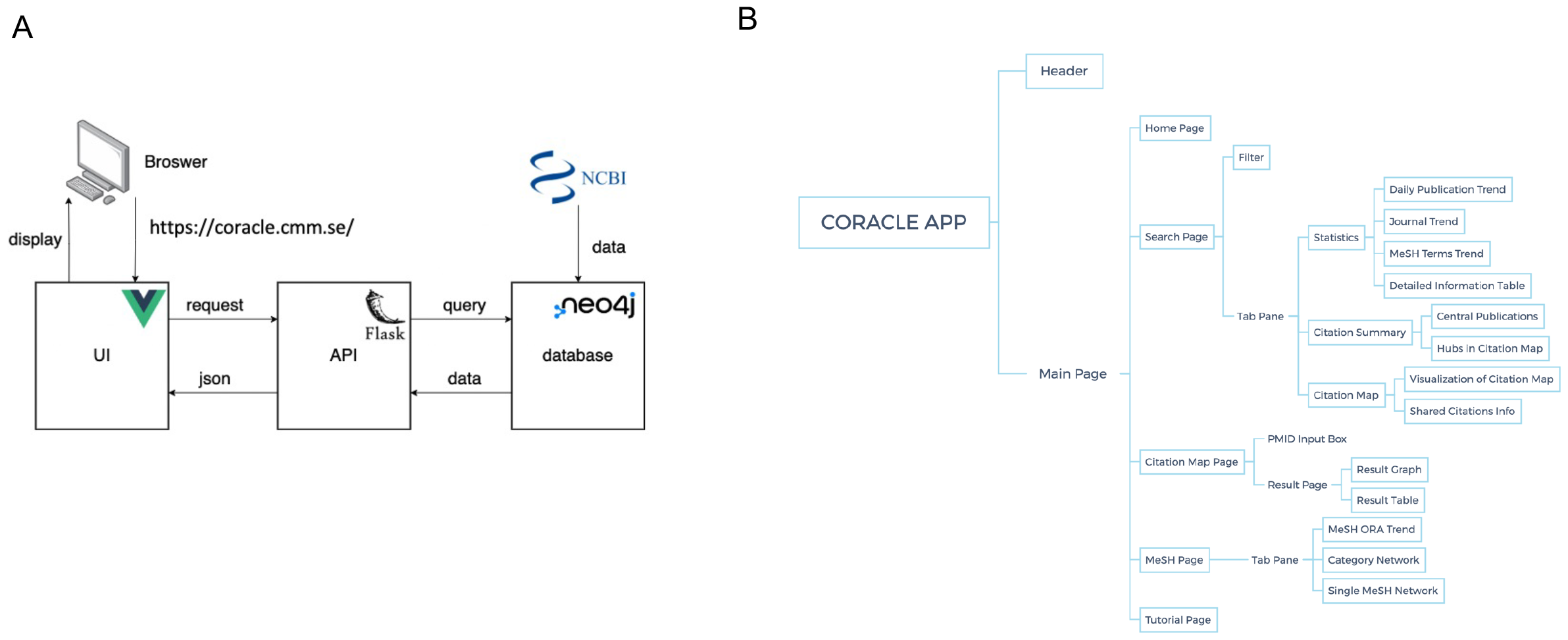
Schematic overview of the CORACLE application. (A) Schematic overview of data processing in CORACLE. (B) An overview of modules currently available in CORACLE. Shapes with borders are components, and shapes without borders are for presenting the structure.

The JavaScript framework Vue.js we utilized for developing user interfaces, using Vite version 2.7.2 for building the Vue components. The structure of the components is shown in Figure 1B. The five main components in the Main Page are controlled by Vue Router version 4.0.12. Components in the Tab Pane are only shown separately in different tabs and they are under the same path. In the Search Page component, Vuex version 4.0.2 is used for passing the change of data in the Filter component to Statistics, Citation Summary, Citation Map components, and their child components conveniently. Some reuse components that do not have specific functions are not shown in the **Figure 1B**, such as the component for drawing loading status.

Flask version 2.0.2, a micro web framework written in Python, is used for building APIs. The extension flask_restful is used in this project to create RESTful APIs since it provides a cleaner way to parse arguments, format output, and organize the routing of calls to the API. After parsing the arguments, the Python script utilizes them to retrieve data from the database. Subsequently, the retrieved data is converted to the JSON format before being sent to Vue components via APIs. The extension flask_cors is employed to handle cross-origin requests, ensuring secure data exchange between the client and server components of the application.

A graph database management system, Neo4j version 4.4.2 is used to save and manage data. Unlike relational databases, graph databases use nodes and edges to store data. To store relationship information, relational databases use tables with one column that contains certain items and another column that contains the related items. For graph databases, relationship information is stored directly as edges. This difference in data structure makes graph databases faster than relational databases when the data is relationships. For the project, different kinds of relationships including the citation relationships and the relationships between articles and Medical Subject Headings (MeSH) terms are used. Therefore, a graph database is suitable for building this application. All information related to articles is fetched from NCBI’s API E-utilities and parsed by the Python module ElementTree, then stored in the Neo4j graph database. Nodes labeled as *COVID* and *Article* are used to save basic article information including PMID, title, journal, publication date, and language. The PMID is saved as a node property called name, while other information is saved as different properties. MeSH and publication types are saved as nodes and connected to COVID nodes since they are in one-to-many relationships with articles. Each reference to an article is created as a node labeled only as *Article* since not all articles in references are COVID-19 related. **Figure 2** shows a scraped article as an example in Neo4j. Due to the data-intense nature and associated long calculation times required for the MeSH functions provided by CORACLE, the data used for the MeSH functions are pre-calculated and stored independently from the data scraped from NCBI E-utility. Given that these relationships do not change markably over time, monthly updates of these terms were deemed sufficient to provide accurate ORA and Fisher’s exact test. To avoid interference with the data directly scraped, a new set of nodes and relationships are created for storing MeSH terms information.

**Figure 2.**
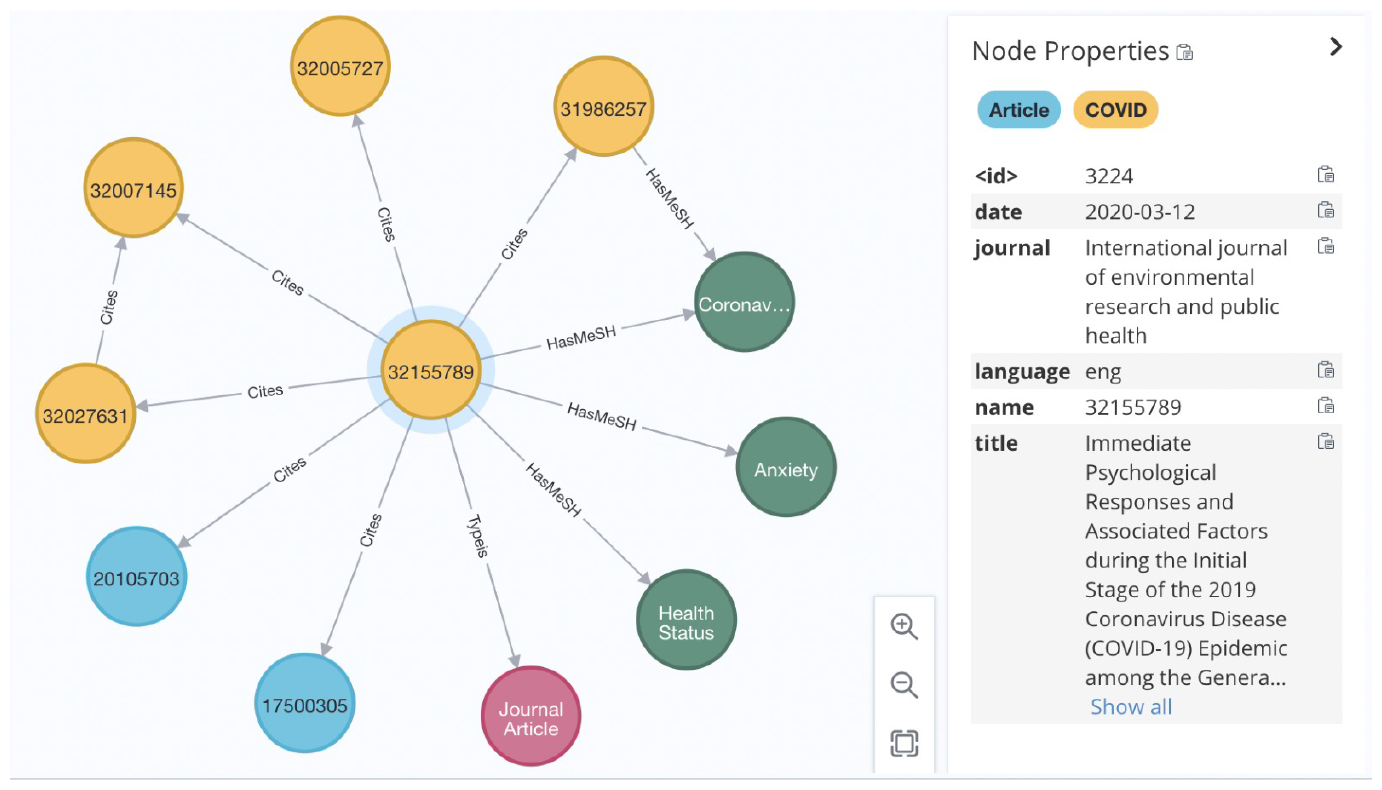
An example of data in Neo4j (not real PMID).

### Functions of CORACLE

In brief, CORACLE is designed to provide 1) fast extraction of emerging hot topics related to SARS-CoV-2/COVID-19, as well as identification of key papers, keywords and journals; 2) personalized extraction of literature of interest by multi-level filters or customized PMID (PubMed Identifier) lists, combined with citation relationships; and 3) help prioritize literature selection and identify highly related publications by direct citation maps and indirect similarity citation networks; 4) understand the relationships between research areas (keywords) 5) generate both human- and machine-readable tables of literature of interest. CORACLE is developed in an Python ^4^ environment, with data mining and integration performed on a once-daily updated literature database from PubMed.

CORACLE includes a global filter block, several visualization panels, and a data table page (see **Figure 3A**, as well as Tutorial). In addition, CORACLE provides a web-based easy-access version with a user-friendly interface. The Search-function consists of five major filter blocks: publication type, country, language, MeSH term, and PMIDs (please see on-line Tutorial for details). The output is generated in four separate blocks (tabs). The STATISTICS block provides summaries of publication trends (by date and cumulatively), publication frequency by journal, as well as a table of the most cited articles. The CITATION MAP block is designed to help identify key publications, i.e., hubs, and publications of putative interest by showing both upstream and downstream citations of the literature of interest.

**Figure 3.**
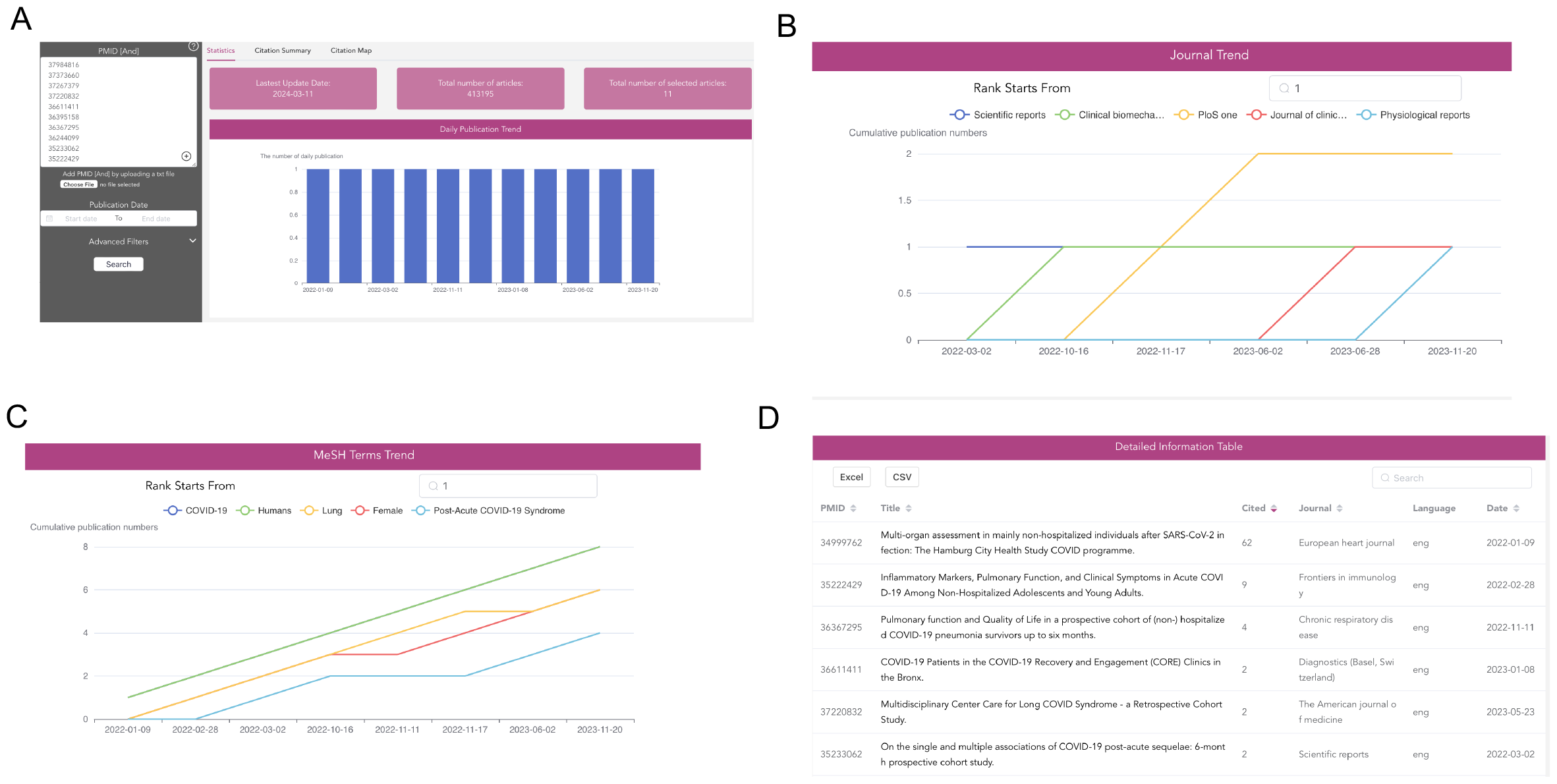
An example of CORACLE utility for literature mining for lung function in patients with post-COVID-19 syndrome. STATISTICS module options. (A) A printscreen from CORACLE showing inputted PMIDs in the left and publication trends on the right. (B,C) Printscreens from the “Statistics” window of CORACLE showing the journal trends and MeSH trends over time. (D) A tabulated summary of the top 6 inputted articles.

The MeSH MAP block outputs network analyses of keyword annotations for publications. The results are displayed as a network map with each MeSH term displayed as a node, and edges representing the number of shared publications between each pair of MeSH terms. All the daily updated background data tables are downloadable for further analysis.

### Literature mining example using CORACLE: post-acute sequele of COVID-19

In the initial stages of the COVID-19 pandemic, healthcare efforts were primarily focused on the immediate recovery of survivors, particularly those suffering from respiratory distress syndrome and COVID-19-related pneumonia. This focus was a direct response to the urgent needs presented by the acute phase of infections. However, as the pandemic progressed, a significant observation was made worldwide. Individuals who had experienced mild COVID-19 symptoms, without requiring hospitalization and showing no abnormalities on X-rays, began to report persistent symptoms^5^. These included heart palpitations, vertigo, brain fog, cough, and breathing difficulties, persisting or recurring weeks to months after initial recovery^6^. Initially, these symptoms were often attributed to the psychological impact of the pandemic, but subsequent reports suggested possible dysregulations in the autonomic nervous system among this patient group^7^.

The persistence of these symptoms, without following any previously known disease patterns, posed significant challenges for healthcare professionals. This was compounded by a lack of published research and clinical guidelines for diagnosing and treating what would eventually be termed Post-Acute Sequelae of SARS-CoV-2 infection (PASC), long COVID, or Post-Acute COVID-19 syndrome (PACS). The condition, characterized by symptoms such as fatigue, shortness of breath, and cognitive dysfunction, impacts daily life significantly. These symptoms can emerge post-recovery or persist from the initial illness, with severity fluctuating over time^8^.

Our clinic has been actively monitoring non-hospitalized PASC patients, focusing particularly on respiratory symptoms and conducting comprehensive investigations. Despite normal spirometry results, radiological findings in some patients have shown air trapping, raising questions about the underlying mechanisms of PASC-associated respiratory issues. The absence of a clear pathophysiological understanding of PASC has made clinical consultations challenging, highlighting the need for systematic, evidence-based approaches.

To bridge the knowledge gap, especially regarding patients with initially mild symptoms, we undertook an extensive literature review. By using findings from our in-house developed system, CORACLE, we aim to provide a comprehensive overview of the publications related to PASC, with a particular focus on the respiratory system. This effort seeks to shed light on the long-term effects of PASC, including the poorly understood phenomena of dysfunctional breathing and its potential long-term impact on lung function.

By design, CORACLE undergoes daily updates to incorporate literature related to COVID-19. Consequently, the default configuration—in the absence of a manually specified list of PMIDs—provides an overview of recent developments within the domain over the preceding months. Between September 14, 2023, and March 14th, 2024 the database has registered 30,129 manuscripts on COVID-19. With winding down of the pandemic, an evident turn towards publications on long COVID, is noticeable. The most frequently cited amongst these is a seminal study on immune profiling in long COVID conducted by Klein et al., within the Mount Sinai-Yale Long COVID project^9^. This investigation undertakes a detailed immune profiling of peripheral blood mononuclear cells (PBMC) from a diverse cohort of 275 individuals, encompassing SARS-CoV-2-infected healthcare personnel, asymptomatic non-infected controls, convalescent controls, subjects with symptoms of long COVID, and individuals with persistent symptoms from a different study. The findings of the study demonstrate an elevated prevalence of non-conventional monocytes (CD14^low^CD16^high^) within the long COVID cohort, alongside a diminished frequency of conventional dendritic cells in comparison with other study groups. Moreover, the research delineated a relative increase in immune reactivity in cells from participants of the long COVID group, as shown by both autoantibody assays against endoproteome and in vitro T-cell stimulation assays. The group also attempted to utilize machine learning approaches to identify the features characteristic for long COVID. In summary, this work - identified as the most cited paper on COVID-19 in the last 6 months by CORACLE – demonstrates that there are immunological changes in the group of long COVID patients. The authors, however, themselves address the point that the groups were formed by the sampling convenience principle, hence the observed heterogenicity. The study was also performed on PBMCs although there are distinct organ-specific symptoms that require elucidation.

For a more targeted search and the demonstration of CORACLE utility, below we outlined example where we aimed to investigate the current state of research regarding to the lung function of individuals experiencing PASC, focusing on those who did not require hospitalization during the primary SARS-CoV-2 infection. As terminology surrounding this phenomenon varies among centers, with the World Health Organization (WHO) and the Centers for Disease Control and Prevention (CDC) predominantly employing the term “post COVID-19 condition.” Within the PubMed database, the MeSH term “Post-Acute COVID-19 syndrome” is designated for this syndrome, however, other variations such as “long COVID-19,” “PASC,” and “long-haul COVID” are commonly used in the literature. Consequently, we attempted to include a comprehensive range of the terms in our search and inputted “Post-acute COVID-19 syndrome” OR “long COVID” OR “PACS” OR “Post-acute sequelae of COVID-19” AND “non-hospitalized” AND “lung function.”

The search for this inquiry yielded a total of 11 publications, with the identified publications compiled into a list of PMIDs and inputted into the CORACLE search interface (**Figure 3A, left panel**)^10–20^. We retained the default settings allowing for broad inclusion without discrimination based on article type, journal, or publication date, however, these advanced filtering options are available for more nuanced analysis.

Subsequently, utilizing CORACLE, we conducted a detailed examination of the interactions among the identified publications concerning lung function in patients with long COVID-19. The platform’s “<u>Statistics</u>” module provided an overview of daily publication trends (**Figure 3A, right panel**), journal distribution (**Figure 3B**), and temporal patterns of MeSH term utilization (**Figure 3C**). Additionally, a comprehensive exportable table-like format facilitated detailed summary of publication statistics, including citation counts, publication details, and access links to PubMed (**Figure 3D**).

Further, the “<u>Citation Summary</u>” feature presented a graphical depiction of citation networks, highlighting “hub” publications and their respective number of citations (**Figure 4A**).

**Figure 4.**
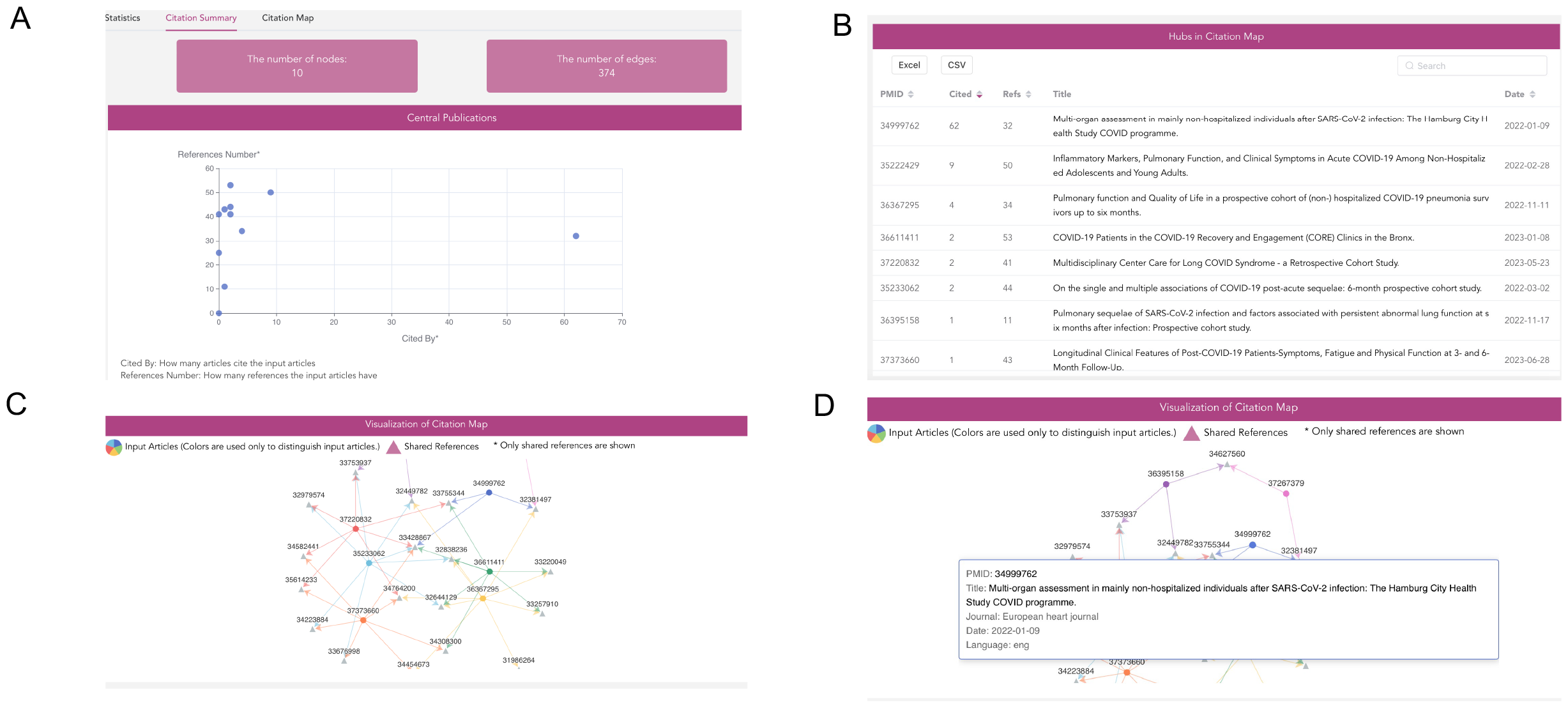
An example of CORACLE utility for literature mining for lung function in patients with post-COVID-19 syndrome. CITATION SUMMARY (A, B) and “CITATION MAP” module options. (A) A printscreen from CORACLE showing the relation between the number of references and number of citations for the inputted publications (B) An exportable summary of all references in the current network (includes both the inputted articles and the articles referenced in them). (C) The citation map for the given search. Dots are hubs, triangles are cited papers. (D) A zoomed-in view on the most cited paper

The “<u>Citation Map</u>” function provided an interactive visualization of citation networks, revealing central nodes within the literature network (**Figure 4C**). Among these was the paper with PMID 34999762 (**Figure 4D**), which gathered considerable citation attention regarding lung function in long-COVID-19 patients^10^. In-depth analysis of individual publications revealed notable insights, such as the study by PMID 33428867, which exhibited considerable citation impact (highest degree) on the papers dealing with lung function in long-COVID-19 patients (**Figure 4C**) ^21^. Degree indicates the number of input articles which either refer to the article in the total section (indegree), or refer to another input article in the search (outdegree). It should, however, be noted that this paper was later retracted and the journal awaits submission of a corrected version.

In this example, the highest-cited publication (**Figure 4A, 4B**) was the paper by Petersen et al. from *Eur Heart J* (with 62 citations) entitled “Multi-organ assessment in mainly non-hospitalized individuals after SARS-CoV-2 infection: The Hamburg City Health Study COVID programme”^10^.This paper examines 443 patients and 1328 matched controls from general population on average 9,6 months after the acute infection. Using body plethysmography (which includes spirometry measurements), the authors identified mild changes in total lung volume and in total airway resistance, as well as a number of mild cardiological changes.

The second most cited paper (a hub with the PMID: 35222429) is by Berven et al^11^. This longitudinal study examines adolescents and young adults aged 12-25 who experienced a mild course of acute COVID-19 infection, compared to age- and sex-matched controls without a recorded history of COVID-19. The study is part of the Long-Term Effects of COVID-19 in Adolescents (LoTECA) project, conducted in two major hospitals in Southeast Norway. The median period between the positive PCR test and measurements was 18 days, establishing baseline data for future observations in this specific age group. Notably, inclusion criteria for the project required confirmation of SARS-CoV-2 infection through a PCR test. In total, the study included 405 cases and 109 matched controls (PCR-negative for SARS-CoV-2 infection, with additional testing for SARS-CoV-2 antibodies).

All included individuals underwent a multiplex cytokine array analysis to measure a panel of inflammatory cytokines, as well as spirometry to measure forced vital capacity (FVC) and forced expiratory volume in one second (FEV_1_). Participants also completed a questionnaire consisting of 24 symptoms rated on a 5-point severity scale.

While significant differences in plasma inflammatory cytokine levels were observed between the COVID-19 and control groups, no significant differences were detected in spirometry measurements. Symptom severity ratings were higher among individuals with detected COVID-19 infection; however, a positive correlation was only observed with female sex and age. Importantly, no correlations were found between pulmonary function parameters and symptom severity in this study.

Another prospective cohort study is presented by de Roos et al.^14^ within the next largest hub in the identified network of publications. In this study, the authors examine the patients 3 and 6 months after the infection and investigate diffusion capacity (DLCO) and quality of life (QoL). Although the total group of included individuals is heterogeneous, the patients are sub-divided according to the severity of the acute infection during the data analysed. All non-hospitalized patients (n=59) – the group we were interested in by limiting our initial literature search - were categorized as moderate cases. 44% of patients in this group had decreased DLCO at 3-months follow up. At 6-month follow up, there was a small yet significant improvement in DLCO for the moderate COVID-19 group, however, it did not normalize neither in severe nor in moderate patient groups at the latest observation time point.

From the “Citation map” block, it is also apparent that the paper with PMID 36611411^20^ entitled “COVID-19 Patients in the COVID-19 Recovery and Engagement (CORE) Clinics in the Bronx” from Diagnostics (Basel) has the highest number of references (**Figure 4B**) and thus can be used for collection of the background articles. This study investigates various clinical parameters, including pulmonary function tests, lung imaging, and symptom assessment, in patients approximately 5 months after experiencing COVID-19 infection. The findings reveal that patients commonly report the onset of multiple new symptoms, such as dyspnea, fatigue, low exercise tolerance, and cognitive difficulties, irrespective of whether they were hospitalized during the acute phase of COVID-19 infection. However, abnormalities in lung function values (including FVC, FEV_1_, DLCO%, and oxygen saturation (SpO_2_) following a 6-minute walking test) were observed exclusively in the group of patients who had been hospitalized during the acute phase. Likewise, the presence of lung opacity persisting after COVID-19 infection was observed solely among patients who had been hospitalized during the acute phase, whereas non-hospitalized patients displayed normal lung imaging findings. Consequently, despite both hospitalized and non-hospitalized patients reporting comparable frequencies and severities of symptoms, lung function abnormalities were exclusively identified in the hospitalized cohort.

Collectively, a brief analysis of the three most cited papers examining lung function in non-hospitalized patients following COVID-19 infection reveals a lack of consistent evidence supporting a uniform decline in lung function across multiple medical centers. Notably, the most cited papers predominantly originate from early studies, published between 2021 and 2022, during the emergence of post-COVID-19 as a distinct medical concern, when its public health implications were not yet fully understood.^21^

This demonstration of literature tracking for lung function in non-hospitalized patients with long COVID-19 exemplifies the utility of CORACLE, which offers versatile applications across various research domains. To the best of our knowledge, our approach to citation map construction, extraction, and analysis represents a novel approach to study and visualize relationship between research articles.

### Building MeSH term networks in CORACLE

An independent computational module designated “MeSH” is available for rapid analysis of interactions among Medical Subject Headings (MeSH) terms in the context of COVID-19. This module by default displays MeSH terms identified through overrepresentation analysis at the time of update. Users can modify term categories, p-value thresholds, time frames, and the volume of analyzed literature (**Figure 5A**).

**Figure 5.**
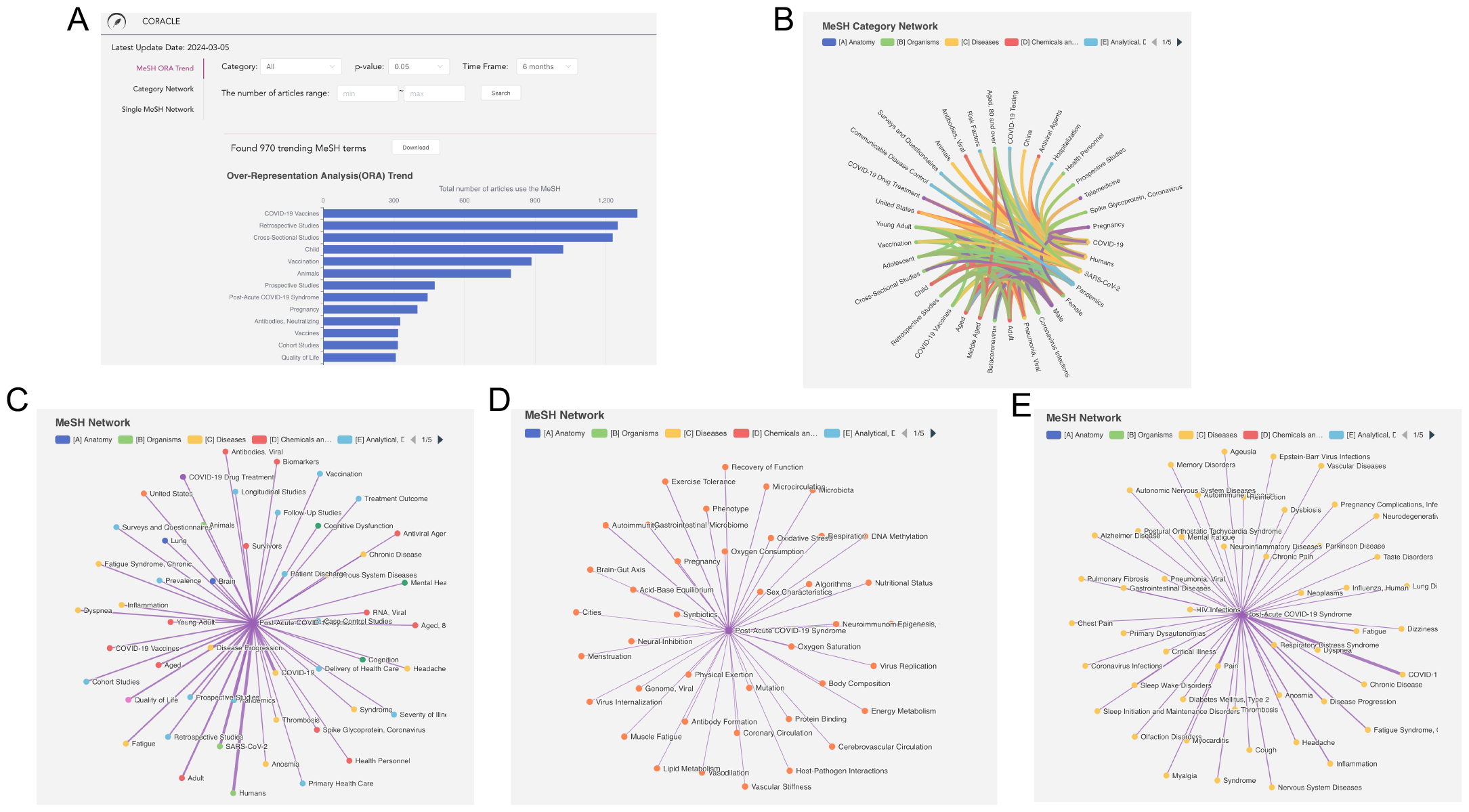
MESH usage in CORACLE (A) MeSH over-representation analysis trends through a default setting. (B) MeSH category network analysis (default, all MeSH terms). (C-E). Single MeSH network for the term “Post-Acute COVID-19 Syndrome” for all terms (C), categorized for “chemicals and drugs” (D) and “Diseases” (E).

In the Category Network Tab, interactions between MeSH terms are depicted via a chord diagram, with correlations determined through pre-calculated Fisher’s exact tests as exemplified on **Figure 5B** with the default settings. The diagram’s line thickness signifies the co-occurrence frequency of term pairs within the literature corpus. Interaction with the diagram, such as hovering over connections, reveals the count of articles for each term pair. By default, connections among the top 300 most prevalent term pairs are shown, though adjustments are possible for both category and minimum article count for displayed terms. Elevating the maximum article count threshold to 4000 allows for the exclusion of commonly occurring terms (e.g., “Humans,” “COVID-19,” “SARS-CoV-2”), enabling a more nuanced examination.

The Single MeSH Network Tab employs a force-directed graph to visualize associations with a selected MeSH term. Initially, this graph displays the 50 most prevalent associated term pairs, with a complete list downloadable in .csv format. The selected input term is marked as a square, with associated terms as circles, and line width reflects co-occurrence frequency. User interaction with the graph, such as hovering to reveal article counts for term pairs, is supported. Adjustments to the minimum article count criteria allow for focused analysis, with a 4000 maximum count limit to exclude prevalent terms for targeted inquiry. **Figure 5C** illustrates the MeSH network for “Post-acute COVID-19 syndrome” across all categories and specifically within “Chemicals and Drugs” (**Figure 5D**) and “Diseases” (**Figure 5E**) categories, offering insight into prevalent diseases and drugs related to “Post-acute COVID-19 syndrome” term usage.

### Limitations with the CORACLE platform

The CORACLE platform, while robust in its design and functionality, exhibits some limitations that need to be considered. Firstly, an important limitation relates to the reporting of references from PubMed. It has been observed that references are not always consistently reported by PubMed, which can result in misleading statistics regarding citation and references, potentially leading to the erroneous assumption that certain articles lack references altogether.

Secondly, the search functionality within the application relies on MeSH terms for article retrieval. While MeSH terms offer standardized indexing of biomedical literature, their utilization often yields broad search results. This broadness becomes particularly apparent in the context of emerging topics such as the Omicron variant and other viral strains like Alpha, Beta, and Epsilon, which lack specific MeSH terms. Consequently, studies focused on these variants may not be effectively captured through MeSH-based searches. Moreover, the classification of various viral strains, including sublineages, under a singular MeSH term such as SARS-CoV-2 further compounds this limitation.

Thirdly, the incompleteness of data poses a significant challenge to the CORACLE platform. As the NCBI’s API E-utility serves as the primary data source, any deficiencies in the initial data retrieval process are inherently unavoidable. A notable shortfall arises in the form of incomplete reference lists within certain articles. Additionally, discrepancies in publication dates further exacerbate data integrity concerns. The adoption of XML-based publication date formats, albeit aimed at standardization, introduces complexities, especially following format alterations within the NCBI’s API. Notably, book-type articles, which are delineated by the absence of the ‘PubMedPubDate’ tag, necessitate alternative methods for date extraction, thereby complicating the data integration process.

Moreover, the manual addition of MeSH terms post-publication introduces another layer of potential data incompleteness. While efforts are made to periodically update and supplement missing information through data retrieval scripts, the efficacy of this approach is contingent upon the availability of updated data within the NCBI databases.

In summary, while CORACLE offers valuable insights into COVID-19 research, these limitations underscore the need for continued refinement and adaptation to ensure the platform effectiveness and relevance in the rapidly evolving landscape of literature related to COVID-19 and post-COVID-19 syndrome.

### Future CORACLE developments and applications

In a rapidly developing research field, keep-up with the scientific literature often represents a bottleneck both for clinicians struggling to provide state-of-the-art treatment to their patients, and pre-clinical researchers aiming to identify novel molecular drug targets and develop novel treatment options. In an emerging, dymanic research field such as that of SARS-CoV-2/COVID-19, both experienced and researchers novel to the field may be limited by the time-consuming task of keeping up with a fast growing body of publications.

Building on our extensive experience of network medicine, we have developed the CORACLE tool to provide a novel means of interactive, multi-level filtering and integration of SARS and COVID literature. The association- and network-based analysis of publications, with MeSH/keywords in the background, represents additional information in the view of data mining. Notably, the multi-level AND and OR combined search of keywords, customized PMID list, and subsequent integration with its summary in publications with time facilitates a time-efficient and comprehensive overview of present and emerging research hotspots. Furthermore, the customized PMID list input search facilitates a personalized prioritization of the citation map output for experts in the field.

In the future, CORACLE expansion directs firstly to incorporate the option of including papers in adjacent fields, such as SARS-CoV-1 and MERS. Then to extract and predict COVID-19-related genes and pathways, which will benefit the development of vaccines and the identification of novel therapeutic targets, primarily to protect sensitive populations prone to develop severe COVID-19 disease, including patients with COPD and severe asthma. The potential to extract putative drug target genes and pathways from the emerging literature against the background of established pathway databases represents another untapped area in data mining ^22–24^.

The daily database updates and extensive personalized prioritization schemes, combined with the planned expansion of the potential to include other conditions, as well as integration with various pathway tools to facilitate the identification of novel putative pharmaceutical targets, holds the promise of revolutionizing literature mining as we know it.

## Author contributions

Conception and design: CXL, ÅMW; analysis and interpretation: CXL, ÅMW; drafting of manuscript: KP, PL, IK, C-X. Li, ÅMW; Python processing of PubMed and LitCovid data: YL; Tutorials and example development: KP, PL, IK, CXJ, JG, ÅMW.

## Acknowledgements

We would like to extend our appreciation to Dr. Zhiyong Lu and Dr. Qingyu Chen from National Center for Biotechnology Information (NCBI), National Library of Medicine (NLM) and National Institutes of Health (NIH) for their supports of the batch download of LitCovid data, and to Marika Ström and Benedikt Zöhrer from Karolinska Institutet for their assistance with beta-testing. The project was supported by the Swedish Heart-Lung Foundation and the Swedish Research Council.

## Notes

https://coracle.cmm.se

### Competing Interest Statement

The authors have declared no competing interest.

